# Additions to the list of arthropods of Reunion Island

**DOI:** 10.1101/2024.09.03.610934

**Authors:** Samuel Nibouche, Joëlle Sadeyen, Sébastien Dervin, Mirana Gauche, Sabine Merion, Janice Minatchy, Romuald Fontaine, Philippe Reynaud, Bruno Michel

## Abstract

**Contribution to the inventory of arthropods of Reunion Island**

This work is a synthesis of new taxa for the Reunion Island identified between 1963 and 2022 and not published to date. The collections were carried out mainly by researchers and technicians from CIRAD or FDGDON. Sampling mainly targeted at crop pests and their natural enemies. The list includes 101 taxa new for Reunion. The Hymenoptera and Hemiptera are the most represented orders.

---

The list of Insecta Linnaeus, 1758 and Arachnida Cuvier, 1812 from Reunion Island (21.12° S, 55.5°E) in 2021 included 3155 and 216 taxa respectively in the French national taxonomic reference system TAXREF (Gargominy *et al*., 2021). Legros *et al*. (2020) estimated, by statistical methods, the expected number of taxa in Reunion (taxa already known and taxa not yet reported) at 7673 for Insecta and 824 for Arachnida. These authors estimate that 62% of the terrestrial arthropod fauna of Reunion remains to be discovered.

The work presented here is a contribution to the inventory of arthropods in Reunion. The list is a synthesis of taxa not published to date, collected between 1963 and 2022 mainly as part of crop pest and beneficial identification activities.

## Materials and methods

### Material examined

The collections were carried out over a period extending from 1963 to 2022. The collections mainly targeted crop pests and beneficials. The specimens were kept dry (before 2015) or in ethanol 95° to -30°C (from 2015). The specimens are stored in the collections of the CBGP (Centre de Biologie pour la Gestion des Populations) in Montpellier (https://doi.org/10.15454/D6XAKL) or in the collection managed by the CIRAD (Centre de Coopération Internationale en Recherche Agronomique pour le Développement) at the Pôle de Protection des Plantes (3P) in Saint-Pierre in Reunion Island. The specimens were morphologically identified by the co-authors of this work or sent to specialists for morphological identification (see details in Results section). Five taxa were identified exclusively by barcoding (see below) using the BIN (Barcode Index Number) assignment tool of the BOLD database (Ratnasingham & Hebert, 2013).

### Sequencing

The specimens collected from 2015 onwards were characterised by barcoding.

DNA extraction from the specimens was performed non-destructively using the Qiagen DNeasy Blood & Tissue Kit (Qiagen, Courtaboeuf, France). DNA samples are stored at 3P.

We sequenced a fragment at position 5’ of the mitochondrial gene for Cytochrome c oxidase I (COI-5P) (Hebert *et al*., 2003). However, in the case of Pseudococcidae, difficulties in PCR amplification led us to amplify instead a portion at the 3’ end of the gene (COI-3P). The positions of the amplified gene portion and the primer pairs used are detailed in Table I, the PCR conditions are indicated by the associated bibliographic references. The sequencing of PCR products was subcontracted to a private company.

**Table I.** Primers used for barcoding.

| Portion of the gene sequenced | Forward Primer | Primer Reverse | Reference | Orders, families |
| --- | --- | --- | --- | --- |
| COI-5P | LCO1490puc_t1<br>LCO1490Hem1_t1<br>LCO1490Hem1_t1 | HCO2198puc_t1<br>HCO2198Hem1_t1<br>HCO2198Hem2_t1 | Germain *et al.* (2013) | Thysanoptera (part), Aleyrodidae (part) |
| COI-5P | LCO1490 | HCO2198 | Folmer *et al.* (1994) | diverse |
| COI-5P | PcoF1 | Lep-R1 | Park *et al.* (2010);<br>Hebert *et al.* (2004) | Monophlebidae |
| COI-3P | C1-J-2183 (aka Jerry) | C1-N-2568 | Simon *et al.* (1994);<br>Palma-Jiménez & Blanco-Meneses (2016) | Pseudococcidae |

**Table II.**
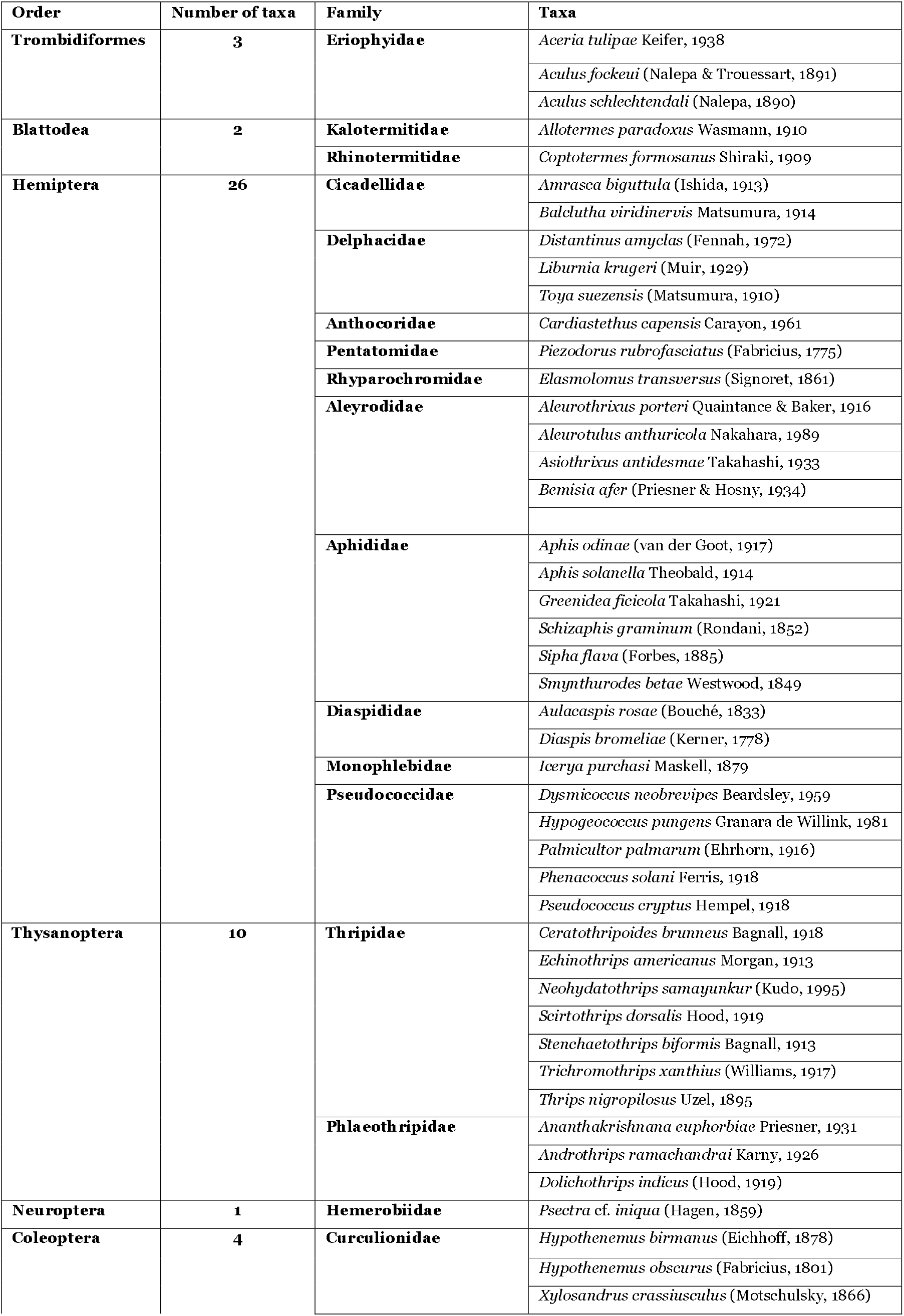
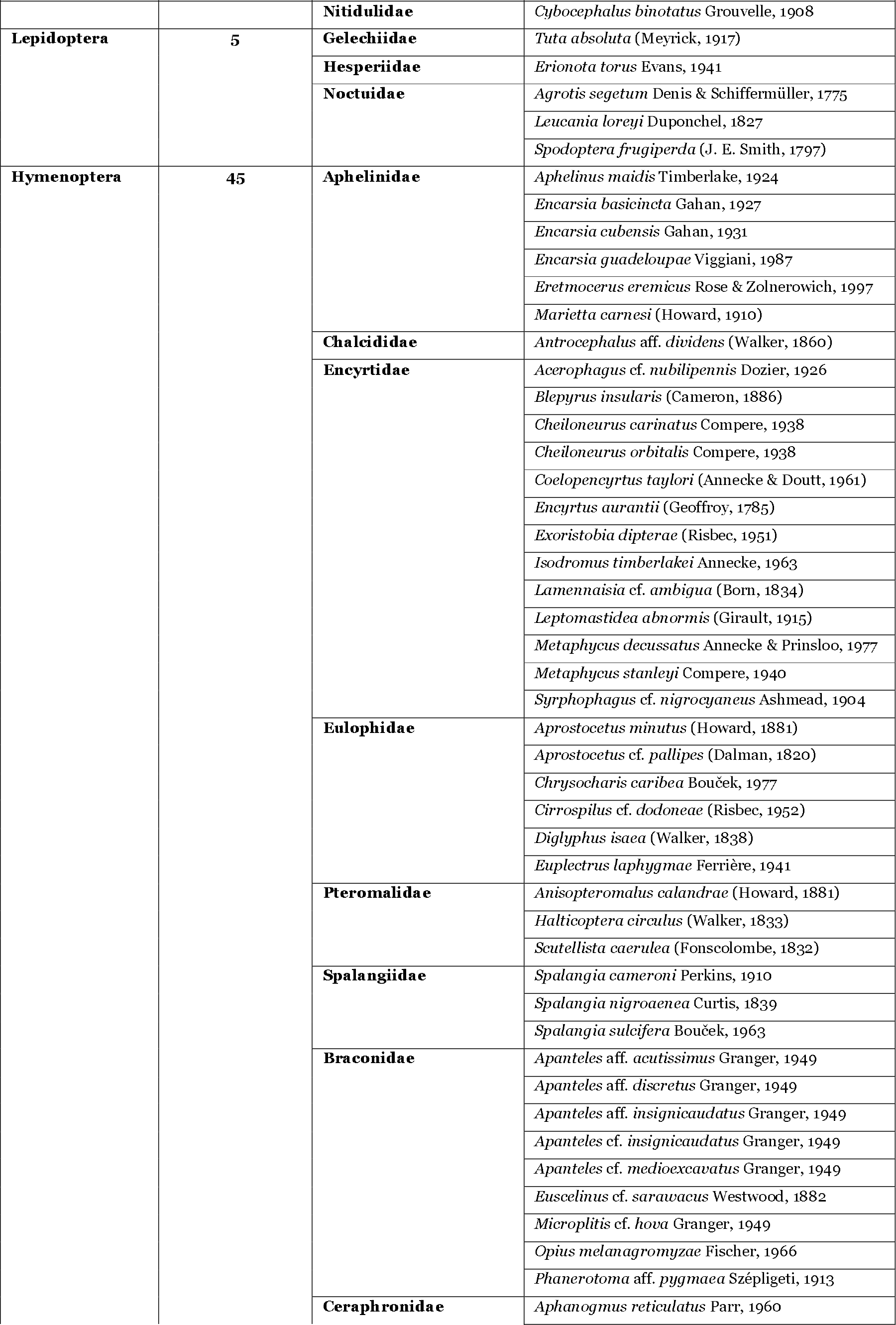
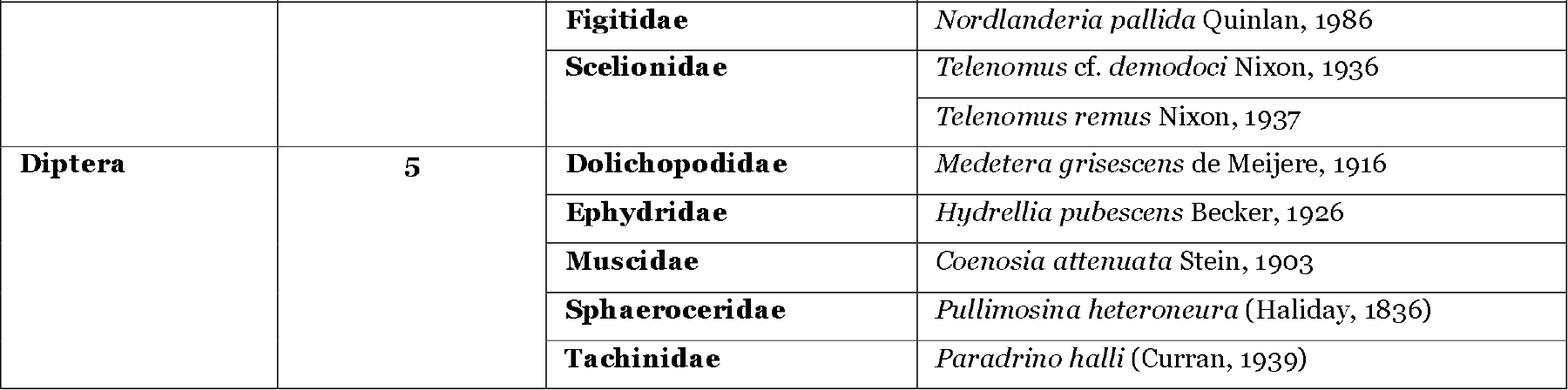
List of 101 taxa new for Reunion Island.

The sequences obtained are available in the Arthemis database (INRAE-CBGP, 2022). They have also been deposited in the BOLD database (Sujeevan & Hebert, 2007), in the DS-UPDRUN1 dataset (dx.doi.org/10.5883/DS-UPDRUN1).

## Results

### List of equipment examined

Specimens with a code in XXXXnnnnn_nn format can be consulted on the Arthemis online database (INRAE-CBGP, 2022), with the exception of those whose code begins with ‘FAUN’. Unless otherwise stated, the specimens collected are imagos. In the absence of any mention of the collector or the number of specimens collected, these are unknown.

Abbreviations and symbols used:

- det.: determinator
- col.: collector
- (num. spec.): numerous specimens collected
- SPV: Plant Protection Service
- LSV: Plant Health Laboratory
- ^#^: barcoded specimen
- ^§^: Taxon identified by barcoding without morphological confirmation

**Arachnida Cuvier, 1812**

**Trombidiformes Reuter, 1909**

**Eriophyidae Nalepa, 1898**

***Aceria tulipae*** Keifer, 1938

Det. Gutierrez Jean-Paul, col. Quilici Serge, La Réunion, 01.VI.1970, *Allium sativum*; det. Gutierrez Jean-Paul, col. Etienne Jean, La Réunion, 01.VI.1970, *Allium cepa*.

***Aculus fockeui*** (Nalepa & Trouessart, 1891)

RQ 470 / FAUN03049, det. Fauvel G., col. Quilici Serge, Saint-Leu, la Chaloupe Saint-Leu, 11.III.1983, *Prunus persica*.

***Aculus schlechtendali*** (Nalepa, 1890)

RQ 475 / FAUN03052, det. Fauvel G., col. Quilici Serge, Saint-Paul, Le Guillaume, 14.III.1983, *Malus pumila*.

**Insecta Linnaeus, 1758**

**Blattodea Brunner von Wattenwyl, 1882**

**Kalotermitidae Froggatt, 1897**

***Allotermes paradoxus*** Wasmann, 1910

RQ 2780 / FAUN12335, det. Ruelle J.E., col. Quilici Serge, La Réunion, Bois Rouge, 03.IV.1992, *Mangifera indica*; RVA 840 / FAUN15340, det. Aberlenc Henri-Pierre, col. Vayssière Jean-François, Saint-Pierre, 27.XII.1997, *Litchi chinensis*.

**Rhinotermitidae Froggatt, 1897**

***Coptotermes formosanus*** Shiraki, 1909

FAUN12262, det. Ruelle J.E., col. Vercambre Bernard, Saint-André, 21.IX.1992; RQ 840 / FAUN04609, det. Ruelle J.E., col. Quilici Serge, Saint-Denis, 01.VIII.1984, *Citrus* sp.; RQ 3330 / FAUN15111, det. Ruelle J.E., col. Quilici Serge, Saint-Louis, Rivière Saint-Louis, 10.I.1997, *Prunus persica*.

**Hemiptera Linnaeus, 1758**

**Auchenorrhyncha Duméril, 1805**

**Cicadellidae Latreille, 1802**

***Amrasca biguttula biguttula*** (Ishida, 1913)

JHUA00001_01 ^#^ (3 specimen), det. Nibouche Samuel, col. Huat Joël, Saint Pierre, Bassin Plat, 04.VI.2020, *Solanum melongena*.

***Balclutha viridinervis*** Matsumura, 1914

MGAU00026_01 (2 specimens), det. Bonfils Jacques, col. Reynaud Bernard, Salazie, Forêt de Bélouve, 28.II.1997.

**Fulgoromorpha Evans, 1946**

**Delphacidae Leach, 1815**

***Distantinus amyclas*** (Fennah, 1972)

RQ 3022 / MGAU00115_01 (1 specimen), det. Bonfils Jacques, col. Quilici Serge, Saint-Philippe, Vallée Heureuse, 30.IV.1995.

***Liburnia krugeri*** (Muir, 1929)

RR 61 / MGAU00129_01 (2 specimens), det. Bonfils Jacques, col. Reynaud Bernard, La Réunion, Ravine Trois Bassins, 05.III.1997, Poaceae.

***Toya suezensis*** (Matsumura, 1910)

RR 67 / MGAU00113_01 (3 specimens), det. Bonfils Jacques, col. Reynaud Bernard, Saint-Paul, Etang Saint-Paul, 05.III.1997, *Paspalidium* sp.

**Heteroptera Latreille, 1810**

**Anthocoridae Fieber, 1836**

***Cardiastethus capensis*** Carayon, 1961

PR 018 / FAUN17085, det. Maldès Jean-Michel, col. Fabre F., La Plaine des Cafres, Piton Hyacinthe, 05.VI.2000, *Brassica oleracea*.

**Pentatomidae Leach, 1815**

***Piezodorus rubrofasciatus*** (Fabricius, 1775)

RV 1952 / MGAU00221_02 (1 specimen), det. Stonedahl G., col. Vercambre Bernard, Saint-Pierre, Bassin Plat, 21.III.1989.

**Rhyparochromidae Amyot & Audinet-Serville, 1843**

***Elasmolomus transversus*** (Signoret, 1861)

RQ 1637 / FAUN07342, det. Slater J.A., col. Quilici Serge, Le Tampon, 19.XI.1987, *Fragaria* sp.

**Sternorrhyncha Duméril, 1806**

**Aleyrodidae Westwood, 1840**

***Aleurothrixus porteri*** Quaintance & Baker, 1916

RQ 1911 / FAUN08501, det. Bink-Moenen R.M., col. Quilici Serge, La Possession, Ilet Latanier, 25.IV.1988, *Citrus latifolia*.

***Aleurotulus anthuricola*** Nakahara, 1989

LSV1200009 / JSTR00070_01 (8 specimens), det. Streito Jean-Claude, col. Minatchy Janice, Sainte-Marie, 29.XII.2011, *Anthurium* sp.

***Asiothrixus antidesmae*** Takahashi, 1933

C17-1231 / LSV1702599, det. Germain Jean-François, col. Vitry Eric, Le Port, 07.XII.2017, *Anthurium* sp.

***Bemisia afer*** (Priesner & Hosny, 1934)

FAUN17032, det. Maldès Jean-Michel, col. Delatte Hélène, Saint-Pierre, Ravine-des-Cabris, 05.IX.2000, *Manihot esculenta*; FAUN17252, det. Delvare Gérard, col. Grondin & Thornary, Sainte-Rose, Piton Sainte-Rose, 17.V.2000, *Manihot esculenta*.

**Aphididae Latreille, 1802**

***Aphis odinae*** (van der Goot, 1917)

RQ 1502 / FAUN06285, det. Leclant François, col. Quilici Serge, Etang-Salé, 30.VII.1986, *Muraya paniculata*; RQ 3202 / FAUN14495, det. Leclant François, col. Quilici Serge, Saint-Benoît, 12.VI.1996, *Litchi chinensis*.

***Aphis solanella*** Theobald, 1914

RVA 829 / FAUN15329, det. Leclant François, col. Vayssières Jean-François, La Plaine des Cafres, Piton Hyacinthe, 02.II.1998, *Solanum nigrum*; RQ 2107 / FAUN08282 (num. spec.) det. Leclant François, col. Manikom Ronald & Quilici Serge, Saint-Denis, La Bretagne, 11.I.1989, *Solanum americanum*.

***Greenidea ficicola*** Takahashi, 1921

C14-242/ LSV1400561 (4 specimens), det. Balmès Valérie, 18.VI.2014, *Ficus* sp.

***Schizaphis graminum*** (Rondani, 1852)

FAUN09408, det. Leclant François, col. Reynaud Bernard, Le Tampon, Bois Court, 15.XI.1989, *Triticum aestivum*.

***Sipha flava*** (Forbes, 1885)

C17-966 / LSV1702154 (36 specimens), det. Mouttet Raphaëlle, col. FDGDON Réunion, Saint-Denis, 09.X.2017, *Saccharum* sp.; RTIB00049_01 ^#^ (num. spec.), det. Nibouche Samuel, col. Tibère Richard, Saint-Denis, La Bretagne, 26.IX.2017, *Saccharum* sp.; RTIB00048_01 ^#^ (num. spec.), det. Nibouche Samuel, col. Tibère Richard, Sainte-Marie, La Mare, 26.IX.2017, *Saccharum* sp.; MDUP00006_01 ^#^ (num. spec.), det. Nibouche Samuel, col. Duployer Marianne, Sainte-Marie, La Mare, 19.X.2017.

***Smynthurodes betae*** Westwood, 1849

Det. Balmès Valérie, col. Lucas Eric, Le Tampon, 05.II.2008, *Solanum lycopersicum*; det. Mintachy Janice, col. Ferrante A., Le Tampon, 30.IV.2008, *Phaseolus vulgaris*; det. Balmès Valérie, col. Mérion Sabine, Cilaos, 30.VII.2008, *Lens culinaris*.

**Diaspididae Targioni-Tozzetti, 1868**

***Aulacaspis rosae*** (Bouché, 1833)

C18-109 / LSV1800284, det. Germain Jean-François, col. Duployer Marianne, Les Avirons, 19.II.2018, *Cycas* sp.

***Diaspis bromeliae*** (Kerner, 1778)

Det. D’Aguilar J., col. Garçonnet C., La Réunion, 01.V.1965, *Ananas comosus*

**Monophlebidae Maskell, 1880**

***Icerya purchasi*** Maskell, 1879

SDER00394_01 ^#^ (num. spec.), det. Nibouche Samuel, col. Dervin Sébastien, Etang-Salé, 10.I.2017, *Sophora tomentosa*; BREY00004_01 (num. spec.), det. Germain Jean-François, col. Reynaud Bernard, Etang-Salé, 03.XI.2017, *Phyllanthus casticum*; MDUP00009_01 ^#^ (num. spec.), det. Nibouche Samuel, col. Duployer Marianne, La Plaine des Cafres, 02.IV.2019, *Acacia* sp.

**Pseudococcidae Heymons, 1915**

***Dysmicoccus neobrevipes*** Beardsley, 1959

LMUL00035_02 ^#^ (7 specimens), det. Germain Jean-François & Nibouche Samuel, col. Muller Lucile, Saint-Pierre, Pierrefonds, 30.I.2015, *Mangifera indica*; MATI00079_01 ^#^ (3 specimens), det. Nibouche Samuel, col. Atiama Morguen, Saint-Gilles, Grand Fond, 26.X.2017, *Mangifera indica*; MATI00080_01 ^#^ (3 specimens), det. Nibouche Samuel, col. Atiama Morguen, Saint-Gilles, Grand Fond, 26.X.2017, *Annona reticulata*; BHOS00004_01 (3 specimens), det. Balmès Valérie, col. Hostachy Bruno, Les Avirons, 2020, *Agave* sp.

***Hypogeococcus pungens*** Granara de Willink, 1981

C19-28 / LSV1900220, det. Balmès Valérie, col. Pallas Régine, Les Avirons, 22.I.2019, *Hylocereus undatus*.

Poveda-Martínez *et al*. (2019, 2020) suggest the existence of cryptic species within *Hypogeococcus pungens populations*. In the current state of the literature, our specimens having been collected on Cactaceae are to be linked to *Hypogeococcus pungens sensu lato*; *Hypogeococcus pungens sensu stricto* corresponding to the populations present on Amaranthaceae.

***Palmicultor palmarum*** (Ehrhorn, 1916)

L2017. RE5.099 / BHOS00001_01 ^#^ (3 specimens), det. Germain Jean-François, col. Hostachy Bruno, Les Avirons, 20.XI.2017, *Dypsis lutescens*.

***Phenacoccus solani*** Ferris, 1918

LSV1102034 / EPIE00394 (1 specimen), det. Germain Jean-François, Saint-Leu, 17.VIII.2011, *Solanum lycopersicum*.

***Pseudococcus cryptus*** Hempel, 1918

LSV1401292 / SDER00205_01 ^#^ (3 specimens), det. Germain Jean-François, La Réunion, 20.VIII.2014, Arecaceae.

**Thysanoptera Haliday, 1836**

**Terebrantia Haliday, 1836**

**Thripidae Stephens, 1829**

***Ceratothripoides brunneus*** Bagnall, 1918

C18-299 / LSV1800588, det. Reynaud Philippe, col. Fontaine Romuald, Saint-Joseph, 12.IV.2018, *Solanum lycopersicum*.

***Echinothrips americanus*** Morgan, 1913

ISA5 / ICAB00001_01 (3 specimens), det. Michel Bruno, col. Cabeu Isabelle, Saint-Pierre, Bassin Martin, 11.XII.2014, *Rosa* sp.; M3 / MTEN00022_01 (1 specimen), det. Michel Bruno, col. Tenailleau Mickael, Saint-Pierre, Bassin Martin, 12.VIII.2014, *Rosa* sp.; SDER00264_01 ^#^ (1 ex.), det. Frago Enric & Nibouche Samuel, col. Frago Enric, La Réunion, Saint-Pierre, Bassin Martin, 22.VIII.2016, *Rosa* sp.; LSV1401764 / SDER00241_01 ^#^ (num. spec.), det. Balmès Valérie & Nibouche Samuel, La Réunion, 28.X.2014, *Rosa* sp.

***Neohydatothrips samayunkur*** (Kudo, 1995)

C18-867 / LSV1801179, det. Reynaud Philippe, col. Fontaine Romuald, Saint-Paul, 17.VII.2018, *Tagetes patula*

***Scirtothrips dorsalis*** Hood, 1919

Det. LSV Montpellier, col. Vitry Eric, Saint-Paul, 07.VI.2012, *Dendrobium* sp.; LSV1500777 / SDER00239_01 (num. spec.), det. Balmès Valérie, La Réunion, 29.VII.2015, *Rosa* sp.

***Stenchaetothrips biformis*** Bagnall, 1913

MJAQ00108_01 (1 specimen), det. Michel Bruno, col. Jacquot Maxime, Saint-Paul, Boucan Canot, 21.VIII.2014; MJAQ00109_01 (1 specimen), det. Michel Bruno, col. Jacquot Maxime, Saint-Pierre, Pierrefond, 20.VIII.2014, *Mangifera indica*.

***Trichromothrips xanthius*** (Williams, 1917)

C17-1230 / LSV1800164, det. Reynaud Philippe, col. Pallas Régine, Salazie, 04.XII.2017, *Anthurium* sp.

***Thrips nigropilosus*** Uzel, 1895

C11-1655 / LSV1500777 (5 specimens), det. Balmès Valérie, col. Pallas Régine, Etang Salé, 27.IX.2011, *Lactuca sativa*.

**Tubulifera Haliday, 1836**

**Phlaeothripidae Uzel, 1895**

***Ananthakrishnana euphorbiae*** Priesner, 1931

C22-01 / LSV2200039 / MGAU04292 ^#^ (25 specimens), det. Reynaud Philippe, col. Danger Camille & Hoarau Henri, Saint-Leu, Pointe au Sel, 10.XII.2021, *Euphorbia puntasalinae* sp. nov.

***Androthrips ramachandrai*** Karny, 1926

Det. LSV Montpellier, col. Bagny Patricia, Saint-Joseph, 04.VII.2012, *Ficus* sp.; LSV1102694 / SDER00233_01 (num. spec.), det. Reynaud Philippe, La Réunion, 09.XI.2011, *Ficus* sp.

***Dolichothrips indicus*** (Hood, 1919)

33BM / SDER00230_01 (6 specimens), det. Michel Bruno, col. Quilici Serge, Saint-Pierre, Bassin Plat, 01.I.1998, *Malus domestica*.

**Neuroptera Linnaeus, 1758**

**Hemerobiidae Latreille, 1802**

***Psectra cf. iniqua*** (Hagen, 1859)

MGAU04166_01 ^#^ (5 specimens), det. Michel Bruno, col. Festin Cyril, Saint-Pierre, Bassin Plat, 24.II.2021, *Abelmoschus esculentus, Portulaca* sp. & *Hubertia* sp. (specimens transmitted by Morguen Atiama from a colony reared by the biofactory ‘Coccinelle’).

**Coleoptera Linnaeus, 1758**

**Curculionidae Latreille, 1802**

***Hypothenemus birmanus*** (Eichhoff, 1878)

RQ 2806 / FAUN12081 (9 specimens), det. Wood S.L., col. Normand Frédéric, Saint-André, Ravine creuse, 15.V.1992, *Litchi chinensis*; RQ 3577 / FAUN15743 (19 specimens), det. Aberlenc Henri-Pierre, col. Dubois B., Saint-Pierre, Pierrefonds, 20.XI.1997, *Vitis vinifera*; RQ 3437 / FAUN15749 (5 specimens), det. Aberlenc Henri-Pierre, col. Dubois B., Saint-Pierre, Bassin Martin, 28.VII.1997, *Vitis vinifera*.

***Hypothenemus obscurus*** (Fabricius, 1801)

RQ 863 / FAUN04701 (10 specimens), det. Wood S.L., col. Quilici Serge, Saint-Paul, Piton Maïdo, 31.XII.1984, *Ulex europaeus*.

***Xylosandrus crassiusculus*** (Motschulsky, 1866)

RQ 3221 / FAUN14370 (16 specimens), det. Wood S., col. Quilici Serge, Saint-Benoît, Bras Madeleine, 12.VI.1996, *Litchi chinensis*; RVA 241 / FAUN14677, det. Delvare Gérard, col. Vayssières Jean-François, Bras-Panon, 23.VII.1997, *Litchi chinensis*; RVA 242 / FAUN14677, det. Delvare Gérard, col. Vayssières Jean-François, Bras-Panon, 23.VII.1997, *Litchi chinensis*; JPOU00099_01 ^#^ (1 specimen), det. Nibouche Samuel, col. Poussereau Jacques, Saint-Benoît, Sainte-Marguerite, Chemin de Ceinture, 13.II.2017; JPOU00105_03 ^#^ (3 specimens), det. Nibouche Samuel, col. Poussereau Jacques, Saint-Pierre, Vieux Domaine de La Ravine des Cabris, 10.XII.2016.

**Nitidulidae Latreille, 1802**

***Cybocephalus binotatus*** Grouvelle, 1908

RQ 1355 / FAUN05623 (15 specimens), det. Endrody-Younga S., col. Guyot J. & Manikom Ronald, Sainte-Clotilde, 24.III.1986, *Nerium oleander*, ex *Pseudaulacaspis pentagona*; RQ 2310 / FAUN09730 (53 specimens), det. Delvare Gérard, col. Quilici Serge & Manikom Ronald, Saint-Denis, 22.XII.1989, *Nerium oleander*, ex *Pseudaulacaspis pentagona*; RQ 2318 / FAUN09823 (4 specimens), det. Delvare Gérard, col. Quilici Serge & Manikom Ronald, Saint-Denis, 21.XII.1989, *Nerium oleander*.

**Lepidoptera Linnaeus, 1758**

**Gelechiidae Stainton, 1854**

***Tuta absoluta*** (Meyrick, 1917)

C18-120 / LSV1800374 (4 specimens), det. Ramel Jean-Marie, col. FDGDON Réunion, Saint-Joseph, 22.III.2018, *Solanum lycopersicum*; MDUP00007_01 ^#^ (3 specimens), det. Nibouche Samuel, col. Duployer Marianne, Saint Joseph, La Crête, 17.V.2018, *Solanum lycopersicum*; AFRA00302_01 ^#^, AFRA00303_01 ^#^ & AFRA00304_01 ^#^ (3 specimens), det. Nibouche Samuel, col. Fontaine Romuald, La Réunion, 28.III.2018, *Solanum lycopersicum*; RTIB00114_01 ^#^ (3 specimens), det. Nibouche Samuel, col. Tibère Richard, Saint-Paul, Bois de Nèfles, 01.XI.2019, *Solanum lycopersicum*.

**Hesperiidae Latreille, 1809**

***Erionota torus*** Evans, 1941

C16-323 / LSV 1600637 (1 specimen), det. Mouttet Raphaëlle, col. FDGDON Réunion, Saint-Paul, 02.VI.2016, *Musa* sp.; AFRA00205_01 ^#^ & AFRA00206_01 ^#^ (2 specimens), det. Nibouche Samuel, col. Fontaine Romuald, La Réunion, 23.VI.2016, *Musa* sp.; RFON00001_01 ^#^ (5 specimens), det. Nibouche Samuel, col. Fontaine Romuald, Saint-Paul, 02.VI.2016; JSAD00009_01 ^#^ (2 specimens), det. Nibouche Samuel, col. Sadeyen Joëlle, Saint-Pierre, 02.IV.2017, *Musa* sp.; JSAD00010_01 ^#^ (2 specimens), det. Nibouche Samuel, col. Sadeyen Joëlle, Saint-Pierre, 29.XI.2017, *Musa* sp.; SDER00259_01 ^#^ (1 specimen), det. Nibouche Samuel, col. Simon P., Saint-Denis, 31.VIII.2016, *Musa* sp.

**Noctuidae Latreille, 1809**

***Agrotis segetum*** (Denis & Schiffermüller, 1775)

LCOS00004_01 ^#^ (3 larvae), det. Nibouche Samuel, col. Costet Laurent, Saint-Louis, Les Makes, 01.XII.2014; ISA13 / ICAB00010_01 ^#^ (2 larvae), det. Nibouche Samuel, col. Cabeu Isabelle, Le Tampon, Grand Tampon les Hauts, 17.III.2015, *Fragaria* sp.

***Leucania loreyi*** (Duponchel, 1827)

RTIB00078_01 ^#^ (2 larvae), det. Nibouche Samuel, col. Tibère Richard, Sainte Suzanne, Bagatelle, 03.IX.2015, *Saccharum* sp.; 07-DJP-P1-SPOFRU-9 / EPAY00065_01 ^#^ (1 legs collected from a glued adult), det. Nibouche Samuel, col. Payet Elisa & Gerville Scholastie, Saint Leu, 13.II.2019, pheromal trapping.

***Spodoptera frugiperda*** (J. E. Smith, 1797)

RFON00006_01 ^#^ (5 specimens), det. Nibouche Samuel, col. Fontaine Romuald, La Réunion, 19.IX.2018; RTIB00092_01 ^#^ (3 specimens), det. Nibouche Samuel, col. Tibère Richard, Saint-Leu, Le Plate, 01.I.2019, *Zea mays*; C18-1232 / LSV1801802 (1 specimen), det. LSV Montpellier, col. Fontaine Romuald, La Réunion, 13.IX.2018, *Zea mays*.

**Hymenoptera Linnaeus, 1758**

**Aphelinidae Thomson, 1876**

***Aphelinus maidis*** Timberlake, 1924

RQ 1792 / FAUN07400, det. Hayat M., col. Quilici Serge, Saint-Louis, Le Gol, 10.XII.1987, *Saccharum* sp., ex Aphididae

***Encarsia basicincta*** Gahan, 1927

70P/ FAUN19352, det. Delvare Gérard, col. Borowiec Nicolas, L’Étang-Salé, 06.VII.2006, *Cocos nucifera*, ex *Aleurotrachelus atratus*; 82P / FAUN19361, det. Delvare Gérard, col. Borowiec Nicolas, L’Étang-Salé, 11.X.2006, *Cocos nucifera*, ex *Aleurotrachelus atratus*; 102P / FAUN19457, det. Delvare Gérard, col. Borowiec Nicolas, Le Tampon, La Chatoire, 02.V.2007, *Dictyosperma album*, ex *Aleurotrachelus atratus*; 103P / FAUN19458, det. Delvare Gérard, col. Borowiec Nicolas, L’Étang-Salé, 02.V.2007, *Cocos nucifera*, ex *Aleurotrachelus atratus*; 104P / FAUN19459, det. Delvare Gérard, col. Borowiec Nicolas, L’Étang-Salé, 20.V.2007, *Cocos nucifera*, ex *Aleurotrachelus aratus*; 105P / FAUN194060, det. Delvare Gérard, col. Borowiec Nicolas, Ravine-des-Cafres, Saint-Pierre, 20.V.2007, *Capsicum frutescens*, ex *Aleurotrachelus trachoides*.

***Encarsia cubensis*** Gahan, 1931

106P / FAUN19461, det. Delvare Gérard, col. Borowiec Nicolas, L’Étang-Salé, 02.V.2007, *Cocos nucifera*, ex *Aleurotrachelus atratus*.

***Encarsia guadeloupae*** Viggiani, 1987

FAUN20168, det. Delvare Gérard, col. Borowiec Nicolas, Le Port, 03.IV.2008, *Pritchardia pacifica*, ex *Aleurodicus dispersus*.

***Eretmocerus eremicus*** Rose & Zolnerowich, 1997

PR0038 / FAUN17367, det. Delvare Gérard, col. Fabre F., Saint-Paul, 01.XI.2000, ex *Trialeurodes vaporariorum*.

***Marietta carnesi*** (Howard, 1910)

RQ 893 / FAUN04677 (2 specimens), det. Hayat M., col. Manikom Ronald, Saint-Leu, Colimaçons, 15.VIII.1984, *Pelargonium*, ex *Pseudaulacaspis pentagona*; RQ 904 / FAUN04812 (1 ex.), det. Hayat M., col. Quilici Serge, Saint-Pierre, Bassin Martin, 28.I.1985, *Citrus sinensis*, ex *Chrysomphalus ficus*; RQ 964 / FAUN05353 (2 specimens), det. Hayat M., col. Quilici Serge, Saint-Pierre, Bassin Martin, 25.IV.1985, *Mangifera indica*, ex *Aulacaspis cinnamomi*; RQ 1380 / FAUN05798 (8 specimens), det. Hayat M., col. Quilici Serge, Saint-Denis, 24.IV.1986, *Nerium oleander*, ex *Pseudaulacaspis pentagona*; RQ 1513 / FAUN06269 (5 specimens), det. Hayat M., col. Quilici Serge, Saint-Pierre, Bassin Martin, 26.II.1986, *Citrus* sp., ex *Pseudaonidia trilobitiformis*; RQ 1515 / FAUN06343 (22 specimens), det. Hayat M., col. Guyot J., Sainte-Clotilde, 24.VI.1986, *Nerium oleander*, ex *Pseudaulacaspis pentagona*.

**Chalcididae Latreille, 1817**

***Antrocephalus aff. dividens*** (Walker, 1860)

RQ 2151 / FAUN08559 (1 specimen), det. Delvare Gérard, col. Quilici Serge, Saint-Pierre, Bassin Martin, 29.V.1989, *Citrus sinensis*.

**Encyrtidae Walker, 1837**

***Acerophagus cf. nubilipennis*** Dozier, 1926

RQ 4827 / FAUN21922 (9 specimens), det. Delvare Gérard, col. Fontaine Olivier, La Possession, 29.X.2010, *Manihot esculenta*, ex *Paracoccus marginatus*.

***Blepyrus insularis*** (Cameron, 1886)

RQ 3704 / FAUN15521, det. Delvare Gérard, col. Quilici Serge, Saint-Leu, Piton-Saint-Leu, 31.III.1998, *Annona squamosa*, ex Pseudococcidae.

***Cheiloneurus carinatus*** Compere, 1938

RQ 1493 / FAUN06258, det. Delvare Gérard, col. Quilici Serge, Saint-Denis, La Bretagne, 19.III.1986, *Dombeya acutangula*, ex Pseudococcidae.

***Cheiloneurus orbitalis*** Compere, 1938

RQ 604 / FAUN03674 (10 specimens), col. Quilici Serge, Saint-Benoît, 21.XI.1983, *Citrus* sp., ex Chrysopidae; RQ 1756 / FAUN07360, det. Delvare Gérard, col. Quilici Serge, Saint-Pierre, Bassin Martin, 15.XI.1987, *Citrus* sp., ex Chrysopidae; RQ 1796 / FAUN07407, det. Delvare Gérard, col. Quilici Serge, Saint-Pierre, Bassin Plat, 14.XII.1987, *Citrus* sp., ex Chrysopidae; RQ 1822 / FAUN07409 (58 specimens), det. Delvare Gérard, col. Quilici Serge, Saint-Pierre, Bassin Plat, 15.I.1988, *Citrus* sp., ex Chrysopidae; RQ 1834 / FAUN07414 (5 specimens), det. Delvare Gérard, col. Quilici Serge, Saint-Pierre, Bassin Plat, 19.I.1988, *Citrus aurantiifolia*, ex Chrysopidae.

***Coelopencyrtus taylori*** (Annecke & Doutt, 1961)

FAUN18012, det. Delvare Gérard, col. Caplong Philippe, Saint-Paul, Savannah, 25.II.2003, ex *Xylocopa fenestrata*.

***Encyrtus aurantii*** (Geoffroy, 1785)

RVA 377 / FAUN14699, det. Delvare Gérard, col. Vayssières Jean-François, Saint-Leu, 10.VII.1997, ex Coccidae.

***Exoristobia dipterae*** (Risbec, 1951)

RQ 2894 / FAUN12534, det. Delvare Gérard, col. Montagneux B., Sainte-Marie, La Ressource, 02.XII.1993, *Solanum melongena*, ex *Neoceratitis cyanescens*; RQ 3103 / FAUN13809, det. Delvare Gérard, col. Quilici Serge, La Réunion, 19.X.1995, ex *Neoceratitis cyanescens*.

***Isodromus timberlakei*** Annecke, 1963

RQ 459 / FAUN03037 (2 specimens), det. Delvare Gérard, col. Quilici Serge, Le Tampon, 17.II.1983, *Citrus* sp., ex Chrysopidae; RQ 604 / FAUN03676 (10 specimens), det. Delvare Gérard, col. Quilici Serge, Saint-Benoît, 21.XI.1983, *Citrus* sp., ex Chrysopidae; RQ 1755 & RQ 1758 / FAUN07359 & FAUN07362, det. Delvare Gérard, col. Quilici Serge, Saint-Pierre, 15.XI.1987, *Citrus* sp., ex Chrysopidae; RQ 1825 / FAUN07413, det. Delvare Gérard, col. Quilici Serge, Saint-Pierre, 15.I.1988, *Citrus* sp., ex Chrysopidae; RQ 1793 / FAUN07403, det. Delvare Gérard, col. Quilici Serge, Saint-Pierre, 14.XII.1987, *Citrus* sp., ex Chrysopidae.

***Lamennaisia cf. ambigua*** (Born, 1834)

RVA 1302 / FAUN16539, det. Delvare Gérard, col. Vayssières Jean-François, La Possession, 10.XII.1999, *Cycas* sp., ex Coccidae.

***Leptomastidea abnormis*** (Girault, 1915)

N° 10 / FAUN14134, det. Delvare Gérard, col. Kreiter P., La Réunion, 20.IX.1996, *Solanum auriculatum*, ex Pseudococcidae.

***Metaphycus decussatus*** Annecke & Prinsloo, 1977

FAUN13381, det. Delvare Gérard, col. Goebel Régis, Saint-Denis, la Bretagne, 15.IX.1994, *Saccharum* sp., ex *Pulvinaria elongata*; FAUN13581, det. Delvare Gérard, col. Goebel Régis, Saint-Benoît, 15.III.1995, ex *Pulvinaria elongata*.

***Metaphycus stanleyi*** Compere, 1940

RQ 942 / FAUN05336, det. Delvare Gérard, col. Garin S., La Saline les Bains, 11.III.1985, *Abelmoschus esculentus*, ex *Parasaissetia nigra*.

***Syrphophagus cf. nigrocyaneus*** Ashmead, 1904

R12 / FAUN16553, det. Delvare Gérard, col. Guilloux Thomas, Le Tampon, Bras-de-Pontho, 19.X.1999, *Brassica oleracea*, ex *Episyrphus* sp.

**Eulophidae Westwood, 1829**

***Aprostocetus minutus*** (Howard, 1881)

FAUN13583, det. Delvare Gérard, col. Goebel Régis, Saint-Benoît, 15.III.1995, ex *Pulvinaria elongata*; RQ 874 / FAUN04660, det. Delvare Gérard, col. Quilici Serge, Saint-Denis, 07.I.1985, *Citrus* sp., ex *Coccus viridis*; RQ 1402 / FAUN05819 (24 specimens), det. Delvare Gérard, col. Quilici Serge, Saint-Denis, 29.V.1985, *Citrus* sp., ex *Coccus viridis*; RQ 400 / FAUN02930, det. Delvare Gérard, col. Quilici Serge, Sainte-Suzanne, 14.X.1982, *Eugenia brasiliensis*; RQ 395 / FAUN02949, det. Delvare Gérard, col. Quilici Serge, Sainte-Suzanne, 14.X.1982, *Citrus clementina*, ex *Icerya seychellarum*.

***Aprostocetus, cf. pallipes*** (Dalman, 1820)

RV 2072 / FAUN09871, det. Delvare Gérard, col. Vercambre Bernard, Saint-Louis, Le Gol, 20.II.1980.

***Chrysocharis caribea*** Bouček, 1977

N° 4 / FAUN12053, det. Delvare Gérard, col. Marinot E., Les Avirons, Le Tévelave, 11.V.1993, ex *Liriomyza* sp.; FAUN09996, col. Bordat Dominique, La Réunion, 15.IX.1989.

***Cirrospilus cf. dodoneae*** (Risbec, 1952)

RQ 3002 / FAUN13571, det. Delvare Gérard, col. Quilici Serge, Saint-Pierre, Bassin-Plat, 20.IV.1995, *Citrus* sp., ex *Phyllocnistis citrella*.

***Diglyphus isaea*** (Walker, 1838)

RVA 1291 / FAUN16528, RVA 1292 / FAUN16529, RVA 1293 / FAUN16530, det. Delvare Gérard, col. Vayssières Jean-François, Salazie, Mare à Vieille Place, 02.XII.1999, *Cucumis melo*, ex *Liriomyza huidobrensis*; PR 006/ FAUN17073, det. Delvare Gérard, col. Fabre F., Saint-Pierre, Bassin Martin, 05.V.2000, *Solanum lycopersicum*, ex *Liriomyza* sp.

***Euplectrus laphygmae*** Ferrière, 1941

RVA 392 / FAUN14713, det. Delvare Gérard, col. Vayssières Jean-François, Saint-Pierre, Bassin Martin, 15.V.1997, *Solanum lycopersicum*, ex *Helicoverpa armigera*.

**Pteromalidae Dalman, 1820**

***Anisopteromalus calandrae*** (Howard, 1881)

RV 272 / FAUN02681, det. Delvare Gérard, col. Vercambre Bernard, Saint-Denis, 15.V.1982, foodstuffs, ex Silvanidae.

***Halticoptera circulus*** (Walker, 1833)

RVA 1289 / FAUN16526, det. Delvare Gérard, col. Vayssières Jean-François, Salazie, Mare à Vieille Place, 02.XII.1999, *Cucurbita maxima*, ex *Liriomyza huidobrensis*; RVA 1290 / FAUN16527, det. Delvare Gérard, col. Vayssières Jean-François, Salazie, Mare à Vieille Place, 02.XII.1999, *Cucurbita pepo*, ex *Liriomyza huidobrensis*; RVA 1292 / FAUN16529, det. Delvare Gérard, col. Vayssières Jean-François, Salazie, Mare à Vieille Place, 02.XII.1999, *Cucumis sativus*, ex *Liriomyza huidobrensis*; FAUN09075, det. Delvare Gérard, col. Bordat Dominique, Saint-Paul, L’Hermitage, 16.XI.1989, *Phaseolus vulgaris*, ex *Ophiomyia* sp.

***Scutellista caerulea*** (Fonscolombe, 1832)

P2 / FAUN21924, det. Delvare Gérard, col. Fontaine Olivier, La Possession, 29.X.2010, ex Coccoidea.

**Spalangiidae Haliday, 1833**

***Spalangia cameroni*** Perkins, 1910

FAUN02398, det. Delvare Gérard, col. Vercambre Bernard, Saint-Paul, Saint-Paul, 12.VI.1980, ex *Physiphora* sp.; FAUN02400, det. Delvare Gérard, col. Vercambre Bernard, Saint-Denis, La Bretagne, 30.VI.1980, ex *Stomoxys niger*; FAUN02401, RV 57, det. Delvare Gérard, col. Vercambre Bernard, Saint-Denis, La Bretagne, 30.VI.1980, ex *Stomoxys niger*; FAUN02412, det. Delvare Gérard, col. Barre, Sainte-Marie, 30.IX.1980, ex *Stomoxys calcitrans*; FAUN02424, det. Delvare Gérard, col. Barre, Saint-Paul, Savannah, 29.X.1980, ex *Musca* sp.; FAUN02425, det. Delvare Gérard, col. Barre, La Réunion, 01.X.1980, ex *Musca* sp.

***Spalangia nigroaenea*** Curtis, 1839

RQ 3105 / FAUN13811 (8 specimens), det. Delvare Gérard, col. Goebel M., Saint-Denis, La Bretagne, 19.X.1995, ex *Neoceratitis cyanescens*; FAUN02398, det. Delvare Gérard, col. Vercambre Bernard, Saint-Paul, 12.VI.1980, ex *Physiphora* sp.; FAUN02402, det. Delvare Gérard, col. Vercambre Bernard, Saint-Paul, Petite France, 30.VI.1980, ex *Stomoxys calcitrans*; FAUN02412, det. Delvare Gérard, col. Barre, Sainte-Marie, 30.IX.1980, ex *Stomoxys calcitrans*; FAUN02424, det. Delvare Gérard, col. Barre, Saint-Paul, Savannah, 29.X.1980, ex *Musca* sp.; FAUN02425, det. Delvare Gérard, col. Barre, La Réunion, 01.X.1980, ex *Musca* sp.; FAUN2426, RV 83, det. Delvare Gérard, col. Barre, Saint-Paul, Petite France, 05.XI.1980, ex *Stomoxys calcitrans*.

***Spalangia sulcifera*** Bouček, 1963

RQ 3104 / FAUN13810, det. Delvare Gérard, col. Montagneux B., Saint-Denis, La Bretagne, 19.X.1995, ex *Neoceratitis cyanescens*; RVA 412 / FAUN14732, det. Delvare Gérard, col. Vayssières Jean-François, Saint-Pierre, Bassin Plat, 15.V.1997, ex Dacini; RV 25 / FAUN01557, det. Delvare Gérard, col. Vercambre Bernard, Saint-Denis, La Bretagne, 15.V.1980, ex *Stomoxys calcitrans*; RVA 1046 / FAUN15632, det. Delvare Gérard, col. Vayssières Jean-François, La Possession, 12.XI.1997, *Citrullus lanatus*, ex *Dacus ciliatus*.

**Ichneumonoidea Latreille, 1802**

**Braconidae Nees, 1811**

***Apanteles aff. acutissimus*** Granger, 1949

FAUN11751, det. Delvare Gérard, col. Vercambre Bernard, Saint-Denis, 04.IV.1997, *Momordica charantia*, ex *Diaphania* sp.; RVA 389 / FAUN14710, det. Delvare Gérard, col. Vayssières Jean-François, Saint-Pierre, Bassin Plat, 21.I.1997, *Cucurbita maxima*, ex *Diaphania indica*; RVA 388 / FAUN14709, det. Delvare Gérard, col. Vayssières Jean-François, Saint-Pierre, Mont-Vert les Bas, 04.IV.1997, *Cucumis sativus*, ex *Diaphania indica*; RVA 391 / FAUN14712, det. Delvare Gérard, col. Vayssières Jean-François, Saint-Pierre, Bassin Martin, 21.VIII.1997, *Citrullus lanatus*, ex *Diaphania indica*; RVA 924 / FAUN15424, det. Delvare Gérard, col. Vayssières Jean-François, La Possession, 22.X.1997, *Cucumis melo*, ex *Diaphania indica*; RVA 925 / FAUN15425, det. Delvare Gérard, col. Vayssières Jean-François, Saint-Paul, Saint-Gilles, 04.XI.1997, *Citrullus lanatus*, ex *Diaphania indica*.

***Apanteles aff. discretus*** Granger, 1949

N°608 / FAUN15163, det. Delvare Gérard, col. Attié Marc, Saint-Leu, La Chaloupe, 03.XII.1995, ex *Ischnurges* sp.

***Apanteles aff. insignicaudatus*** Granger, 1949

RV 7 / FAUN01555, det. Delvare Gérard, col. Vercambre Bernard, Petite Île, Piton Goyave, 03.XI.1974, *Cynara cardunculus*, ex *Tebenna micalis*; RVA 709 / FAUN15209, det. Delvare Gérard, col. Vayssières Jean-François, Saint-Pierre, Mont-Vert les Hauts, 18.XI.1997, *Cynara cardunculus*, ex *Tebenna micalis*; RVA 717 / FAUN15217, det. Delvare Gérard, col. Vayssières Jean-François, La Plaine des Cafres, Piton Hyacinthe, 15.XII.1997, *Cynara scolymus*, ex *Tebenna micalis*.

***Apanteles cf. insignicaudatus*** Granger, 1949

R 1 / FAUN16542, det. Delvare Gérard, col. Guilloux Thomas, Le Tampon, Bras de Pontho, 02.XI.1999, *Brassica oleracea*, ex *Plutella xylostella*.

***Apanteles cf. medioexcavatus*** Granger, 1949

RVA 478 / FAUN14768, det. Delvare Gérard, col. Vayssières Jean-François, Saint-Pierre, Ravine des Cabris, 21.I.1997, *Momordica charantia*, ex Lepidoptera; PR 025 / FAUN17092, det. Delvare Gérard, col. Fabre F., Saint-Pierre, Ravine des Cabris, 11.I.2000, *Cucumis melo*, ex *Diaphania indica*; PR 013 / FAUN17081, det. Delvare Gérard, col. Fabre F., Saint-Leu, 20.IV.2000, *Cucurbita* sp., ex *Diaphania indica*.

***Euscelinus cf. sarawacus*** Westwood, 1882

RVA 413 / FAUN14733, det. van Achterberg K., col. Vayssières Jean-François, Saint-Benoît, Sainte-Anne, 05.XII.1997, *Litchi chinensis*, ex Cerambycidae.

***Microplitis cf. hova*** Granger, 1949

RVA 717 / FAUN15217, det. Delvare Gérard, col. Vayssière Jean-François, La Plaine des Cafres, Piton Hyacinthe, 15.XII.1997, *Cynara scolymus*, ex *Tebenna micalis*.

***Opius melanagromyzae*** Fischer, 1966

C22-753B / LSV2200599, det. Rousse Pascal, col. Tisserand Gaëlle, La Réunion, 16.V.2022, *Vigna unguiculata* ssp. *sesquipedalis*, ex Agromyzidae.

***Phanerotoma aff. pygmaea*** Széligeti, 1913

FAUN04667, det. Delvare Gérard, col. Quilici Serge, Saint-Pierre, Bassin Plat, 28.XII.1984, *Citrus aurantiifolia*.

**Ceraphronidea Haliday, 1833**

**Ceraphronidae Haliday, 1833**

***Aphanogmus reticulatus*** Parr, 1960

FAUN13377, det. Delvare Gérard, col. Goebel Régis, Saint-André, 06.VII.1994, *Saccharum* sp., ex *Cotesia* sp.

**Cynipoidea Latreille, 1802**

**Figitidae Thomson, 1862**

***Nordlanderia pallida*** Quinlan, 1986

RVA 521 / FAUN14788, det. Delvare Gérard, col. Vayssières Jean-François, Saint-Denis, 20.VIII.1997, *Allium cepa*, ex *Liriomyza* sp.; RVA 528 / FAUN14793, det. Delvare Gérard, col. Vayssières Jean-François, Saint-André, 02.IX.1997, *Brassica rapa*, ex *Liriomyza* sp.; RVA 552 / FAUN14801, det. Delvare Gérard, col. Vayssières Jean-François, Saint-Denis, 25.IX.1997, *Brassica rapa*, ex *Liriomyza* sp.

**Platygastroidea Haliday, 1833**

**Scelionidae Haliday, 1839**

***Telenomus cf. demodoci*** Nixon, 1936

FAUN18011, det. Delvare Gérard, col. Attié Marc and Morel Gilles, Saint-Denis, 14.III.2003, *Duranta erecta*, ex *Acherontia atropos*.

***Telenomus remus*** Nixon, 1937

FAUN13629, det. Delvare Gérard, col. Goebel Régis, Saint-Denis, 15.IV.1995, ex *Spodoptera mauritia mauritia*; FAUN14026, RG-R, det. Delvare Gérard, col. Goebel Régis, Saint-Denis, 15.IV.1996, ex *Spodoptera mauritia mauritia*.

**Diptera Linnaeus, 1758**

**Dolichopodidae Latreille, 1809**

***Medetera grisescens*** de Meijere, 1916 ^§^

MAJQ00145_01 ^#^ (1 specimen), det. Nibouche Samuel, col. Jacquot Maxime, La Réunion, 20.VIII.2014; MAJQ00149_01 ^#^ (1 specimen), det. Nibouche Samuel, col. Jacquot Maxime, La Réunion, 29.VIII.2013.

**Ephydridae Zetterstedt, 1837**

***Hydrellia pubescens*** Becker, 1926

RQ 4150 / FAUN16923 (2 specimens), det. Martinez M., col. Quilici Serge, Salazie, 14.II.2000, *Nasturtium officinale*; RQ 4151 / FAUN16924 (8 specimens), det. Martinez M., col. Quilici Serge, Salazie, 13.IV.2000, *Nasturtium officinale*.

**Muscidae Latreille, 1802**

***Coenosia attenuata*** Stein, 1903

SDER00147_01 ^#^ (2 specimens), det. Nibouche Samuel, col. Dervin Sébastien, Saint-Pierre, Bassin Martin, 10.V.2015, *Solanum melongena*.

**Sphaeroceridae Macquart, 1835**

***Pullimosina heteroneura*** (Haliday, 1836)

RV 256 / FAUN02695 (1 specimen), det. Deeming J.C., col. Vercambre Bernard, Saint-Joseph, Grand Coude, 10.X.1982, *Phaseolus vulgaris*.

**Tachinidae Fleming, 1821**

***Paradrino halli*** (Curran, 1939)

RV 10 / FAUN01559, det. D’Aguilar, col. Vercambre Bernard, Sainte-Marie, 26.X.1979, *Saccharum* sp., ex *Leucania pseudoloreyi*.

## Discussion

The list includes 101 taxa new to Reunion Island. Five of these taxa were identified using the barcoding technique. Hymenoptera and Hemiptera are the most represented orders, with 45 and 27 taxa respectively. Several invasive pest species of major economic or environmental importance and of recent introduction were identified between 2012 and 2020, such as *Spodoptera frugiperda, Tuta absoluta, Scirtothrips dorsalis, Echinothrips americanus, Erionota torus, Icerya purchasi, Amrasca biguttula biguttula or Sipha flava.* These introductions illustrate the vulnerability of Reunion Island to biological invasions and the current rate of introduction of exotic insects to the island.

## Acknowledgments

This work received financial support from the Epibio project, co-financed by the European Union (ERDF Interreg V), the Reunion region and CIRAD. The authors would like to thank Thibault Ramage for his critical review of the list and Gérard Delvare for his contribution to the Hymenoptera list. Specimens collected after 2019 have been subject to a declaration for access to genetic resources and the sharing of benefits arising from their utilization, within the framework of the Convention on Biological Diversity and the Nagoya Protocol (reference ABSCH-IRCC-FR-248772-1).

